# Beyond pulling: microtubule pushing forces contribute to robust spindle orientation in regular and irregular cell shapes

**DOI:** 10.1101/2025.09.22.677921

**Authors:** Alikhan Yeltokov, Tara M. Finegan, Alexander G. Fletcher, Holly E. Lovegrove, Dan T. Bergstralh

## Abstract

Oriented cell division is fundamental to development and tissue organization, requiring precise control of both spindle positioning and orientation. While cortical pulling forces mediated by dynein motor proteins are well-established drivers of spindle dynamics, the contribution of microtubule polymerization-based pushing forces remains unclear. We developed a generalizable computational biophysical model that integrates both pulling and pushing mechanisms to investigate spindle behavior across diverse cell types and geometries. This model successfully recapitulates experimental observations in three well-studied models: *Drosophila* follicular epithelial cells, *Drosophila* neuroblasts, and the early *C. elegans* embryo. Systematic analysis reveals that while pulling forces are the primary drivers of directed spindle orientation, pushing forces play crucial supporting roles by preventing spindle stalling and promoting alignment dynamics, particularly at high initial misalignment angles. We further applied our model to irregularly shaped zebrafish endothelial cells, which present unique challenges due to their non-spherical morphology and dynamic shape changes during mitosis. Our results demonstrate that asymmetric cortical force generator distributions, potentially localized at cell-cell junctions, can account for the observed off-center spindle positioning in these cells. This work provides a unified framework for understanding how the interplay between cell geometry, molecular polarity cues, and competing physical forces determines spindle dynamics, offering new insights into both canonical and non-canonical division orientations across cell types.

## INTRODUCTION

Oriented cell division is fundamental to the development of all living organisms as it is directly involved in tissue formation and cell fate decisions (reviewed in (Bergstralh et al. 2017)). During this process, chromosomes are separated and pulled toward opposite poles of the cell by the mitotic spindle. This spindle is a dynamic apparatus composed of centrosomes (or spindle poles), microtubules (MTs), and associated proteins (reviewed in (Di Pietro et al. 2016)). MTs are thin, semi-flexible filaments that exhibit dynamic instability—stochastic transitions between growth (polymerization) and shrinkage (depolymerization) phases, driven by the addition or loss of tubulin subunits (Mitchison and Kirschner 1984). In most animal cells, MTs radiate from centrosomes and are organized into three functionally distinct classes: (1) kinetochore MTs, which attach to chromosomes; (2) astral MTs, which extend toward the cell cortex; and (3) interpolar MTs, which overlap with counterparts from the opposite pole (Tolić 2018). Astral MTs grow in a radial, star-like pattern (hence the name ‘astral’) and play key roles in spindle positioning and orientation (reviewed in (Lechler and Mapelli 2021)).

During cell division, the mitotic spindle dynamically adjusts both its orientation and position to ensure proper daughter cell fate and tissue organization. Spindle angle—its orientation relative to the broader tissue architecture - determines the initial placement of daughter cells within the tissue (Figure 1A) (reviewed in (Morin and Bellaïche 2011)). Spindle position—its placement within the cell—determines the division plane and regulates daughter cell size symmetry (Figure 1B), ranging from central (as in most epithelial divisions) to off-center (as in the first division of the *C. elegans* zygote and in zebrafish angiogenesis). Rotational and translational movements of the spindle often occur simultaneously and are driven by forces generated by interactions between astral MTs and the cell cortex (Di Pietro et al. 2016). Together, they precisely orchestrate the cell- and tissue-specific positioning of the division plane, ensuring that daughter cells are born in the correct location, with appropriate size, and equipped with the correct signaling and fate determinants. Misalignment or misplacement of the spindle can have severe consequences, such as disrupting the layered architecture of epithelial tissues or impairing the asymmetric partitioning of fate determinants. Accordingly, improper spindle movements have been linked to tumor formation and developmental disorders such as microcephaly (Fleming et al. 2009; Byrd et al. 2016; Gai and Di Cunto 2017; Quyn et al. 2010; Thoma et al. 2009).

**Figure 1:**
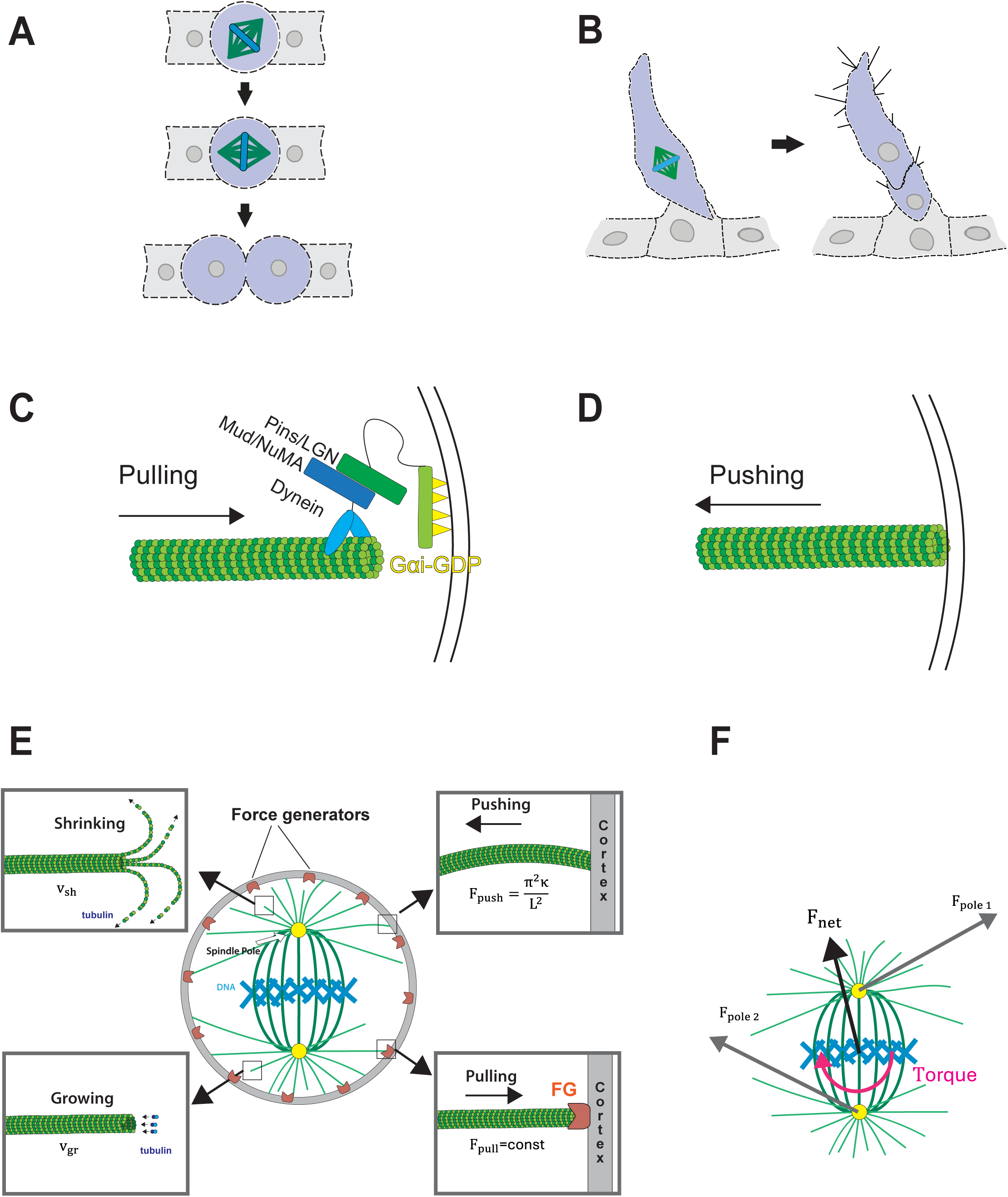
Schematics of spindle orientation and positioning and overview of the computational model. **(A)** Spindle orientation: spindle angle determines daughter cell placement within the tissue in epithelia. **(B)** Spindle positioning: spindle position specifies the division plane and thereby the relative size of the daughter cells in endothelia. **(C, D)** Mechanisms of force generation. **(C)** The canonical spindle orientation machinery, where cortical anchor Gαi, adapter proteins Pins/LGN and Mud/NuMA, and dynein motors generate pulling forces on astral MT ends to align the spindle. **(D)** Growing astral MTs that push against the cortex generate mechanical forces, which are transmitted to the spindle and move it in the opposite direction. **(E)** Schematic summarizing the forces and MT behaviors incorporated in the computational model. Astral MTs undergo stochastic switching between growth and shrinkage. Pushing forces are modeled using Euler beam buckling, while canonical cortical force generators (Gαi–Pins–Mud–dynein) are represented as a single aggregated pulling unit. Force generators are localized to cortical regions defined by polarity cues and exert a constant pulling force (see Methods for details). **(F)** Diagram of force and torque calculations. Forces transmitted by individual MTs to their spindle poles are summed to obtain the net force and torque acting on the spindle, which is assumed to rotate around its center.

Two mechanisms have emerged as potential drivers of spindle orientation and positioning, raising questions about whether and to what extent each works across cell types. The first mechanism involves cortical pulling forces generated by an evolutionarily conserved molecular machinery.

Studies across multiple systems have shown that spindle orientation is typically governed by a suite of at least three proteins: LGN, NuMA, and Gαi in vertebrates; Pins, Mud, and Gαi in *Drosophila*; and GPR1/2, LIN-5, and GOA-1/GPA-16 in nematodes (Lechler and Mapelli 2021). These proteins anchor the MT motor dynein to specific regions of the cell cortex. This localized molecular machinery exerts pulling forces on astral MTs, moving the spindle into the correct position and angle (Figure 1C) (Grill and Hyman 2005; reviewed in (Bergstralh et al. 2017).

The second mechanism involves forces generated through MT polymerization (Figure 1D) (Howard 2006; Howard and Garzon-Coral 2017). When growing MT plus-ends encounter physical barriers, the continued addition of tubulin subunits generates pushing forces - up to 4 pN per MT *in vitro* (Dogterom and Yurke 1997). In a cell with one microtubule-organizing center (MTOC), these pushing forces can collectively guide the MTOC toward the cell’s volumetric center (Holy et al. 1997; Meaders et al. 2020). The importance of pushing has been demonstrated in several systems. In interphase cells, growing MTs that contact the cell cortex can exert forces that displace the nucleus away from the point of contact (Tran et al. 2001; Zhao et al. 2012).

The relative importance of pushing versus pulling mechanisms remains unclear, particularly across different cell types and geometries. While protein knockdown experiments and laser ablation studies have provided valuable insights, they have not conclusively determined the relative contributions of these forces, particularly in cells with diverse geometries and molecular compositions. Additionally, most studies examine spindle positioning and orientation as separate processes, despite their temporal overlap *in vivo*. Experimental models are frequently optimized to investigate one process at the expense of the other—for example, *Drosophila* neuroblasts primarily allow the study of orientation (Cabernard and Doe 2009), as the spindle is spatially constrained and effectively pinned at the cell center, while in the *C. elegans* embryonic zygote, the spindle is already aligned along the division axis, allowing for the study of positioning (Pécréaux et al. 2016). This separation is artificial, as both processes likely involve similar force-generating mechanisms and often occur simultaneously.

Computational modeling offers a powerful approach to address these limitations and test the contributions of different force mechanisms. Previous computational models have been used to study and predict spindle dynamics (Garzon-Coral et al. 2016; Campàs and Sens 2006; Wu et al. 2024; Saleh et al. 2023; Théry et al. 2007; Kozlowski et al. 2007; Kotak et al. 2012). These, and *in vitro* work, have generated compelling evidence for the presence of both pulling and pushing forces acting on mitotic spindles. However, while the role of pulling forces in orienting and positioning mitotic spindles has been studied extensively, the contribution of pushing forces remains unclear.

Current modeling approaches are also limited by their focus on specific cell types and regular geometries. Most existing work examines regularly shaped cells (Théry et al. 2007; Fink et al. 2011), neglecting the morphological diversity increasingly revealed *in vivo* (Lovegrove et al, 2025; Alhashem et al, 2022). Some models have been developed within the context of specific cell types, focusing on fixed geometries and force distributions characteristic of particular biological systems (Nestor-Bergmann et al. 2019; Li and Jiang 2018; Li et al. 2019; Wu et al. 2011; Letort et al. 2016; Kimura and Onami 2005; Wu et al. 2024). To address these challenges, we developed a generalizable computational biophysical model capable of testing pulling and pushing mechanisms across different cell types, including those with dynamic geometries. This model provides a theoretical framework for predicting spindle behavior under different force regimes and in varying cell shapes.

Our results reveal that pushing forces play important but context-dependent roles in spindle dynamics, working synergistically with pulling mechanisms to ensure robust spindle orientation and positioning across diverse cell types and geometries.

## RESULTS

### A novel computational model for spindle orientation

Our model operates in a 2D midplane section of the cell, capturing the azimuthal symmetry of cortical motor proteins (Pins–Mud–Gαi and dynein). This allows for accurate implementation of cortical motor proteins and allows for direct comparison with experimental data. To capture only the essential contributors, we restrict the model to astral MTs, which interact with the cell cortex and strongly influence spindle behavior, while excluding kinetochore MTs that primarily attach to chromosomes. Similarly, we model the canonical pulling machinery (Gαi–Pins/LGN–Mud/NuMA– dynein) as a single aggregated entity, a cortical force generator (FG). The agent-based approach reduces the system to a small set of interacting components—the spindle, MTs, cortical FGs, and the cell cortex (Figure 1E). This representation is at once simple and powerful: instead of tracking all molecular detail, the model highlights how interactions between these agents give rise to emergent spindle behaviors, and how variation in those interactions impacts spindle orientation and positioning.

Astral MTs were treated as straight beams nucleating from spindle poles within a defined fixed angular spread (β angle). They undergo dynamic instability, stochastically switching between growth and shrinkage, with rescue and catastrophe events explicitly modeled. This makes the system inherently noisy, since each MT can behave differently even under identical conditions. When a growing MT reaches the cortex, two outcomes are possible: (1) if it lies within the interaction range of an FG, it binds with a defined probability, and the FG exerts a constant pulling force along the MT axis; or (2) if no FG is nearby, the MT continues to grow against the cortex and generates a compressive pushing force, modeled using Euler buckling mechanics. Each FG was assigned the pulling strength of a single dynein motor, and their localization patterns were adapted from experimentally measured distributions in real cells. The contributions of all individual MTs—whether pulling or pushing—are then summed to determine the total force and torque acting on the spindle at each point in time (Figure 1F).

The position and angle of the spindle evolve based on the total force acting on the spindle. As the spindle changes its position and angle, the astral MTs readjust as well. This can lead to changes in the existing forces or the emergence of new ones. The spindle position and angle were iteratively updated from the current MT configuration, with this feedback repeated in discrete short time steps across the model. Each simulation duration was fixed, based on estimates of observed prometaphase duration derived from previous studies (Bergstralh et al. 2015; Finegan et al. 2019; Neville et al. 2023; Pécréaux et al. 2016). Due to the model’s stochasticity, individual runs can exhibit variability. Therefore, we ran multiple simulations to quantify uncertainty in predictions. Detailed descriptions of the model and its parameters are provided in the Methods, and all parameter values are listed in Supplemental Table 1.

### The model recapitulates spindle orientation in *Drosophila* cell types

As we developed our computational model, we tested whether it could recapitulate observations made in different model systems. We used the *Drosophila* follicular epithelium (FE), a simple monolayer, as a baseline. Mitotic follicle cells tend to orient their spindles along the tissue plane, such that the daughter cells expand the layer (Fernández-Miñán et al. 2007; Bergstralh et al. 2013) (Figure 2A). An advantage to this system is that the location of both Pins and Mud along the mitotic cell cortex has been quantified on the basis of fluorescent signal intensity (Neville et al. 2023). Mud is concentrated on the lateral cell surfaces, in a gradient that is strongest towards the apical cell-cell junction (Figure 2A). We used this gradient as the basis for force generator density in the simulations because Mud, unlike Pins, binds dynein directly. We constructed an angular density distribution by mapping peak Mud signal intensity across all angles, assuming that higher Mud intensity corresponds to a higher local density of cortical force generators.

**Figure 2:**
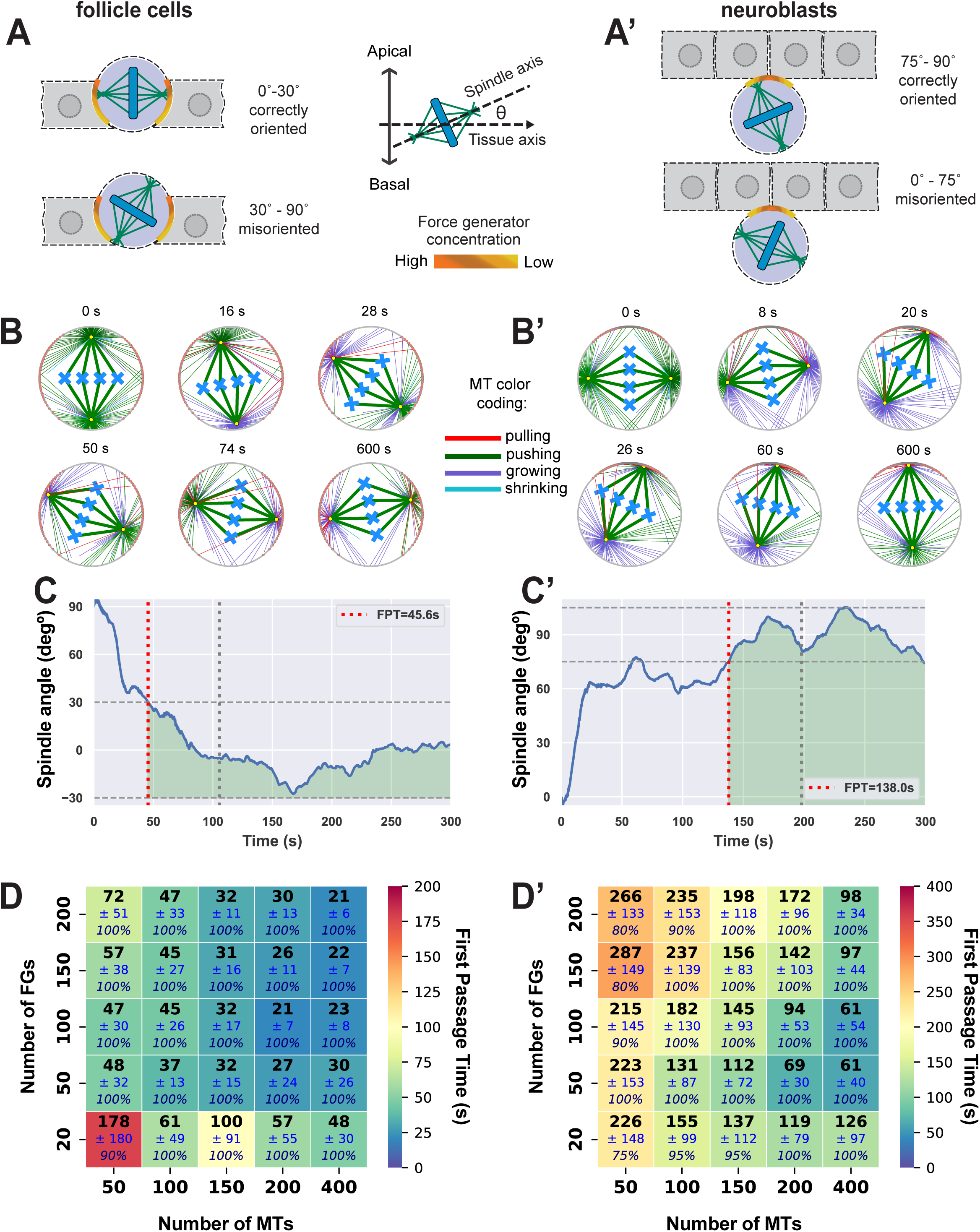
Spindle orientation in flies is recapitulated by the computational model. **(A, A’)** Schematic illustrating the spindle angle measurement convention, defined as the angle between the spindle axis and the tissue polarity axis. In follicle cells **(A)**, a spindle is considered correctly oriented when its angle lies within 0–30° of the tissue axis (horizontal). In neuroblasts **(A’)**, the criterion for correct orientation is 75–90° (vertical). Orange crescents represent cortical force generator (FG) localization, with darker gradients indicating higher density. **(B, B’)** Snapshots of representative simulation runs at successive time steps (Movies S1 and S2). The cell cortex is shown in grey, FGs in orange, centrosomes in yellow, and DNA in cyan. Astral MTs are color-coded as green for pushing, red for pulling, cyan for shrinking, and purple for growing. Initial spindle angles were set to 90° in follicle cells and 0° in neuroblasts. The total number of MTs (*N*_*MT*_) is 200 and the number of FGs (*N*_*FG*_) is 100. **(C, C’)** Spindle angles recapitulate correct orientation for the simulations in **(B, B’)**. Over time, spindle angles converge toward the expected alignment. First passage time (FPT) was used to determine successful orientation, defined as the spindle remaining within the correct angle range for at least 60 seconds. First passage times are 45.6 s for the follicle cell run **(C)** and 138 s for the neuroblast run **(C’)**. Red and grey dotted horizontal lines mark the angle thresholds, while the green shaded region highlights the interval where the spindle is within the accepted range. **(D, D’)** Orientation speed depends on the number of MTs (*N*_*MT*_) and FGs (*N*_*FG*_). For each combination of *N*_*MT*_ and *N*_*FG*_, an average FPT (over 20 simulation runs) is shown in bold black, with standard deviation in navy and percent of successful orientations in italics.

In addition to follicle epithelial cells, spindle orientation has long been investigated in the *Drosophila* neuroblast, a progenitor cell that divides asymmetrically to produce a self-renewing neuroblast and a differentiating cell (Rachel Kraut et al. 1996; Siller and Doe 2009). Cell fate is governed by the distribution of factors including Prospero, Numb, and Brat, that localize to a basal crescent prior to division (Spana et al. 1995; Rhyu et al. 1994; Lee et al. 2006). The spindle-orienting machinery localizes to an apical crescent so that the spindle is oriented along the apical-basal polarity axis, ensuring that only the basal daughter inherits the basally localized factors (Siller et al. 2006). Based on published micrographs (*e.g.* (R Kraut et al. 1996; Wodarz et al. 1999; 2000; Yu et al. 2000; Schaefer et al. 2000; Izumi et al. 2006; Bowman et al. 2006)) we assumed that this crescent is a gradient of force generator density that is strongest in the center (Figure 2A’). A non-uniform, gradient-like distribution of force generators on the cortex provides spatial cues that define a preferred locus—typically the maximum of the gradient—toward which spindle poles are pulled. Like most animal cells, follicle cells and neuroblasts round up to divide (Taubenberger et al. 2020). They are therefore represented simply as circles in our 2D model. We consider spindle length to be equal to the 80% of cell diameter (*L*_*spindle*_ = 0.8 *L*_*cell*_) based on the observed proximity of spindle poles to the cortex (Bergstralh et al. 2013). The initial spindle orientation was set perpendicular to the correct alignment axis: 90° away from the target angle in both follicular epithelium and neuroblasts. For follicular epithelium, this corresponded to a 90° starting angle relative to the horizontal axis, while in neuroblasts it corresponded to a 0° starting angle (Figure 2B, B’ and Movies S1 and S2).

Given the model’s stochasticity, individual trajectories vary, requiring a robust metric to assess successful alignment across many simulations. We defined correct alignment as the spindle reaching the angular range observed for each system (±15° for neuroblasts, ±30° for follicular epithelium) from the corresponding plane (90° for neuroblasts, 0° for follicular epithelium) (Siller and Doe 2008; Bergstralh et al. 2013). To estimate how quickly alignment takes place, we measured the first passage time (FPT), defined as the earliest time the spindle entered this range (Figure 2C, C’). To avoid counting brief random alignments, we required the spindle to remain in the correct region for at least 60 s.

To summarize results across parameter sets, we used a heatmap that displays three metrics for each condition: the mean FPT, its standard deviation, and the success rate (fraction of simulations that achieved and maintained alignment) (Figure 2D, D’). This approach evaluates not only whether alignment occurs, but also how consistently and efficiently it is achieved—providing a comprehensive view of alignment dynamics in a noisy biological system. In addition to FPT, we assessed spindle angle stability after alignment by calculating the post-FPT angle standard deviation (Follicle Cells: Supplemental Figure 1A; Neuroblasts: Supplemental Figure 1B).

Over the numbers simulated we found that astral MT number and force generator number correlate inversely, but imperfectly, with FPT. As the total number of astral MTs and force generators increases, the time it takes for the spindle to orient decreases, though not with a simple, linear relationship. This is likely due to the complexity of the underlying stochastic processes - the force generators require MTs to capture, while free MT plus-ends cause length-dependent pushing. Notably, spindle orientation is less efficient in simulated neuroblasts than in simulated follicle cells. This is likely attributable to the stricter tolerance and the fact that force generators in neuroblasts are positioned in such a way that they can act on only one spindle pole at a time. For the sake of comparison, we also simulated a uniform distribution of FGs in neuroblasts. As expected, the lack of a positional bias decreased the speed of spindle orientation (Supplemental Figure 1A-C). We observed very little impact on final spindle angle (Supplemental Figure 1B).

### The model recapitulates observations in the *C. elegans* embryo

We next challenged our model with a system that presents additional complexity, namely the asymmetrically-dividing *C. elegans* single-cell embryo. The process of spindle orientation in this cell begins with positioning of the two pronuclei. The male pronucleus moves to encounter the female pronucleus, after which both move to the cell center. We simulated the movement of two pronuclei towards the cell center, aligning along the anterior-posterior (A-P) axis (Figure 3A). The pronuclear cortex starts at 80% of the A-P length in the posterior region, with centrosomes aligned orthogonally to the A-P axis. The distribution of force generators is based on published micrographs (Gotta et al. 2003; Srinivasan et al. 2003; Park and Rose 2008). A Gaussian pattern was used to capture enrichment of FGs at both poles, with density decreasing away from the poles, and with the anterior pole having a higher overall FG density than the posterior (3:2 ratio of FG number). Previous modeling work has tested various mechanisms for pronuclear centering, including cytoplasmic pulling (Kimura and Onami 2005), cortical pulling, and a combination of cortical pulling with pushing forces (Coffman et al. 2016). These models explore how different force interactions contribute to the precise alignment and positioning of the pronuclei during cell division. Laser ablations of astral MTs have shown that correct pronucleus complex (PNC) migration required cortical pulling (Wu et al. 2024). The simulations with cortical pulling and pushing reproduced gradual rotation and centering of the PNC (Figure 3A’-A’’, Supplemental Figure 2A, Movie S3), consistent with the asymmetric distribution of force generators, with higher density at the anterior and lower density at the posterior cortex.

**Figure 3.**
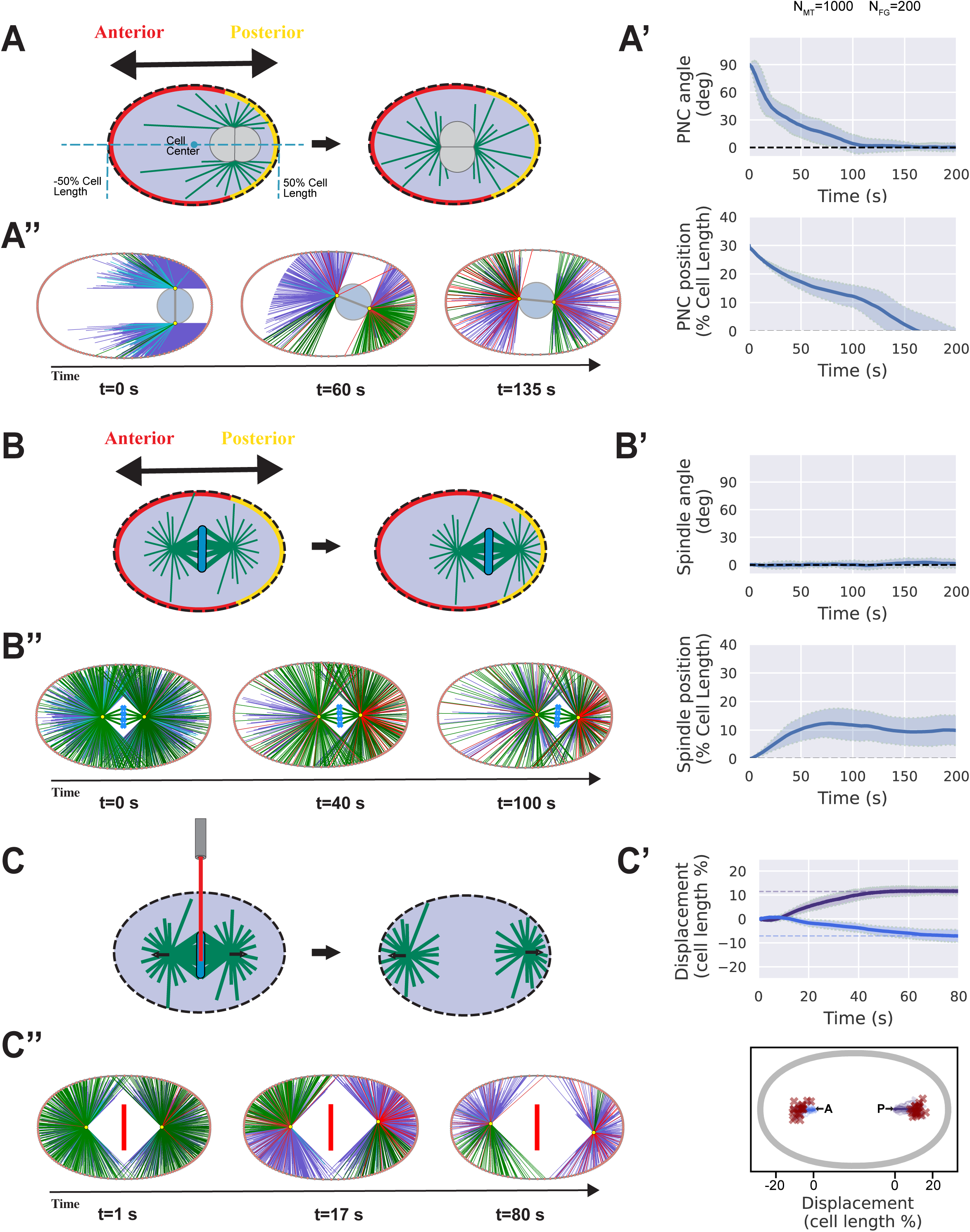
Computational model validation in *C. elegans*. **(A)** Pronuclear complex (PNC) migration in the *C. elegans* embryo. The PNC begins in the posterior one-third of the cell, rotates by 90° to align with the anterior–posterior (A–P) axis, and migrates to the cell center. Anterior and posterior cortical domains are indicated in yellow and red, respectively. **(A’)** Time series of PNC angle (top) and position (bottom) from simulations with *N*_*MT*_ =1000 and *N*_*FG*_ = 200. Blue lines represent the mean of n = 20 runs; shaded regions show standard deviation. **(A’’)** Representative simulation snapshots illustrating sequential stages of migration (Movie S3). Astral MTs are color-coded as in Figure 2. **(B)** Posterior displacement of the mitotic spindle in the *C. elegans* embryo. The spindle is initially centered and subsequently shifts toward the posterior cortex along the A– P axis. Anterior and posterior cortical domains are shown in yellow and red. **(B’)** Time series of spindle angle (top) and position (bottom) for simulations with *N*_*MT*_ =1000 and *N*_*FG*_ = 200. Blue lines indicate the mean of n = 20 runs; shaded regions show standard deviation. **(B’’)** Sequential simulation snapshots illustrating spindle posterior displacement (Movie S4). Astral MTs are color-coded as in Figure 2. **(C)** Simulated spindle severing experiment. A metaphase spindle (L = 22 μm), initially centered, was cut at t = 0 s. The anterior and posterior poles moved apart toward their respective cortical domains. **(C’)** Top: Position versus time plots for posterior (purple) and anterior (blue) poles. Solid lines represent the mean of n = 20 runs; shaded regions indicate standard deviation; dashed horizontal lines are initial positions for both poles, and dashed horizontal lines are final displacements. Bottom: Scatterplot of spindle pole trajectories (color coded as before) and final positions (red X marks). **(C’’)** Sequential snapshots from a representative simulation showing pole separation (Movie S5).

Following this, we simulated spindle positioning after PNC centering and nuclear envelope breakdown. Starting from the cell center, the spindle shifted asymmetrically toward the posterior, consistent with the establishment of posterior displacement in the embryo (Figure 3B). Our model does not incorporate spindle elongation, a process known to be critical for accurate posterior positioning. Nonetheless, these results demonstrate that, even in the absence of elongation, asymmetry in cortical pulling between anterior and posterior domains is sufficient to move the spindle off-center (while maintaining correct orientation) to support asymmetric division (Figure 3B’-B’’, Supplemental Figure 2B, Movie S4).

Foundational evidence for the pulling model is derived from manipulations that disrupt the spindle midzone in the *C. elegans* embryo (Grill et al. 2001) (Figure 3C). In these experiments, movement of the spindle pole(s) away from the impacted site demonstrates that poles are under a cortex-directed pulling force. We simulated ablation of the spindle midzone and found that the spindle poles retract towards the cortex (Figure 3C’-C”, Movie S5), providing further validation of our model. We repeated these simulations using weaker (or absent) pushing forces and observed that, as expected, these parameters cause the poles to move even closer to the cortex (Supplemental Figure 2C, Movie S6).

Taken together, these results indicate that the computational model is generalizable; using parameters derived from the literature it can reproduce experimental observations across different systems.

### A role for MT pushing in spindle orientation

Having validated the model, we next used it to address the question of what role MT pushing performs in spindle orientation. This problem has proven difficult to address experimentally; manipulations aimed at disruption of MT plus-ends also impact MT pulling, and evidence across systems suggests that pulling is the predominant mechanism (reviewed in (Bergstralh et al. 2017)). Spindle orientation generally fails after genetic disruption of the canonical spindle-pulling machinery and in the *C. elegans* embryo, positioning also relies on these factors (Colombo et al. 2003). Polymerization-based pushing can center asters/nuclei and help align spindles, as shown in fission yeast and *Xenopus laevis* (Tran et al. 2001; Sulerud et al. 2020). These findings establish that pushing is a genuine force-generating mechanism, capable of moving large cellular structures and contributing directly to spatial organization.

Our model incorporates these two opposing forces and can therefore be used to explore the contribution of each. We simulated spindle orientation in follicle cells under three conditions: 1) pulling and pushing; 2) pushing only; and 3) pulling only. For the sake of computational efficiency, we used 200 MTs and 100 force generators. These simulations were performed over a range of initial angles from 90° to 45° relative to the ideal final angle (0°) (Figure 4A and Supplemental Figure 3A). In all cases, the pulling and pushing model was most efficient in reaching the final angle (Figure 4A, Supplemental Figure 3A, Movie S1). The pulling-only model lagged, and this effect was most pronounced at the higher initial angles (Figure 4A and Supplemental Figure 3A). The pushing-only model failed to orient the spindle (Figure 4A and Supplemental Figure 3A). Given that pushing forces are isotropic in this cell type, this result is expected and provides further validation of the model.

**Figure 4:**
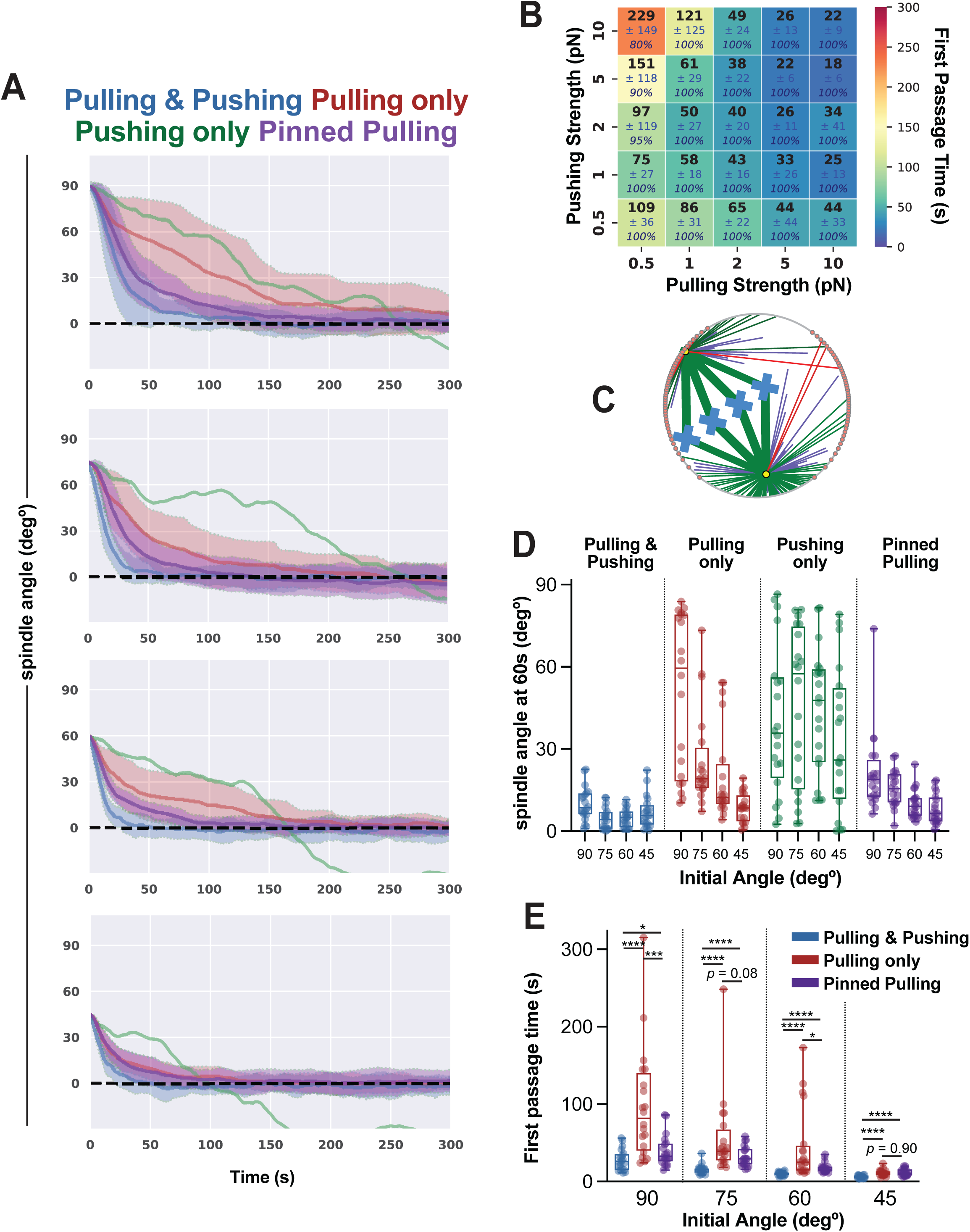
Pushing prevents spindle stalling and accelerate orientation. **(A)** Spindle angle over time in simulated follicle cells. Initial spindle angles were 90°, 75°, 60°, and 45° from top to bottom. Four conditions are shown: combined pulling and pushing (blue), pulling only (red), pushing only (green), and pinned pulling only (purple). Solid lines represent the mean of n = 20 simulations, and shaded envelopes indicate the standard deviation. **(B)** Dependence of spindle orientation speed on MT pulling and pushing strength in the combined condition (*N*_*MT*_ =200 and *N*_*FG*_ = 100). For each parameter combination, the average first passage time (FPT, n = 20 runs) is plotted in bold black, with standard deviation in navy and percent of successful orientations in italics. **(C)** A representative timepoint of a “stalled” spindle. Astral MTs are color coded as in Figure 2B. **(D)** Spindle angle at 60 s across all four conditions and all initial spindle angles. Colors match those in **(A)**. Boxplots display the 25th, 50th, and 75th percentiles, with whiskers spanning the full data range (n = 20). **(E)** Comparison of FPT across combined pulling and pushing, pulling only, and pinned pulling conditions for initial angles of 90°, 75°, 60°, and 45°. Boxplots as in **(C)**. Statistical significance was assessed with the Wilcoxon signed-rank test; *p < 0.05, **p < 0.01, ***p < 0.001, ****p < 0.0001.

As a complementary approach we calculated mean FPTs (Follicle Cells: Figure 4B; Neuroblasts: Supplemental Figure 3B) and post-FPT stabilities (Follicle Cells: Supplemental Figure 3C; Neuroblasts: Supplemental Figure 3D) for simulations over a range of pushing and pulling strengths. These experiments use an initial angle of 90° to maximize the difference across conditions. As expected, mean first passage time and pulling strength are inversely correlated. Remarkably, the same is true for mean passage time and pushing strength. This result suggests that the increased speed of alignment observed in the pushing and pulling model compared to pulling alone (Figure 4A) is due to both forces acting in complementary ways: pulling provides a polarity-dependent bias toward the correct orientation, while pushing, although isotropic and not inherently directional, promotes spindle movement by driving random spins. These spins increase the likelihood that spindle poles come into proximity with cortical regions enriched in force generators, thereby enabling pulling forces to engage more effectively and accelerating overall alignment.

We next addressed the question of why pushing contributes to spindle alignment in these cells. We found that some spindles in the pulling only model stalled out, meaning that they remained stuck at an angle above 45° for an extended period (Figure 4C, Movie S7). This is partly illustrated by the distribution of spindle angles at 60 seconds (simulated time) (Figure 4D); a proportion of spindle angles are observed at or near the initial angle when that angle is 60° or higher. This effect is most obvious at the 90° initial angle condition; at 60 seconds, 8 of 20 spindle angles are clustered around 80° (Figure 4C).

When stalled, one spindle pole becomes closely associated with the cortex while the other pole is pulled away from force generators on the opposing side (Figure 4D). A straightforward explanation is that pushing MTs provide a buffer that prevents translational movement of the spindle. Because the pushing force is inversely proportional to astral MT length, this effect is strongest when MTs are short, allowing the system to sense both spindle position and cell shape. When this buffer is lost, the net force on the spindle depends entirely on pulling forces and their cortical position. Thus, pushing not only accelerates spindle orientation but also acts as a safeguard, preventing the spindle from approaching the cortex too closely and becoming stalled. Stalling would be more likely at higher spindle angles because the concentration of force generators in follicle cells increases towards the apical cell corners (as shown in Figure 2A). To test the importance of buffering, we modeled a condition in which the spindle center is pinned to the cell center, preventing translational motion. (Pushing is not implemented in this condition). “Pinned pulling” provided an improvement in spindle orientation efficiency in comparison to pulling alone, but this effect was only particulary significant at an initial angle of 90° (Figures 4A,E). At the lowest initial angle (45°), there is no difference between the pulling only and pinned pulling models (Figures 4A,E). Together, these results support an important buffering role for the pushing forces generated by growing astral MTs.

Notably, a significant pushing-dependent improvement in spindle orientation efficiency is observed even at lower initial spindle angles (Figures 4A,E), indicating that buffering is not the only contribution of MT pushing. We speculate that pushing MTs also promote dynamicity, and in simulated follicle cells this improves efficiency. Taken together, these results indicate that pushing MTs promote spindle orientation, even in regularly-shaped, symmetrically-dividing cells.

### Vascular endothelial cells are a challenge to convention

Organized growth of the intersegmental vessels (ISVs) in zebrafish larvae occurs through a combination of collective cell migration and cell division. As part of this process, angiogenic endothelial cells take on hierarchical identities as “tip” (leader, at the distal end of new vessel) and “stalk” (follower, the proximal end) cells (Herbert and Stainier 2011). To maintain this hierarchy, tip cells divide asymmetrically to generate a larger (more motile) tip cell and smaller (less motile) stalk cell (Costa et al. 2016; Lovegrove et al. 2025) (Figure 5A). Several features make this system particularly challenging for the study of spindle orientation and position. First, these cells do not round up to divide but rather maintain elongated irregular shapes during mitosis (Figure 5A). Second, to generate asymmetrically sized daughter cells the spindle is offset from the cell center at metaphase (Figure 5A,D,F). Third, the shape of the cell can change over the course of mitosis (Figure 5A). Cell to cell variability in shape, and changes to shape over time, are two features that present a particular challenge to the more conventional modeling approaches used for regularly-shaped cells (as above). We accounted for these by capturing *in vivo* cell outlines over time and translating these outlines into the model (Supplemental Figure 4A). We note that time steps are not identical for each cell but range from 21-60 s per frame. This is a consequence of varied cell geometry; the frame rate was determined by the number of focal places required to capture each cell volume in its entirety. Finally, while the zebrafish is well-suited for studying development through live imaging, genetic techniques to test the roles of pulling and pushing forces have proven challenging.

**Figure 5:**
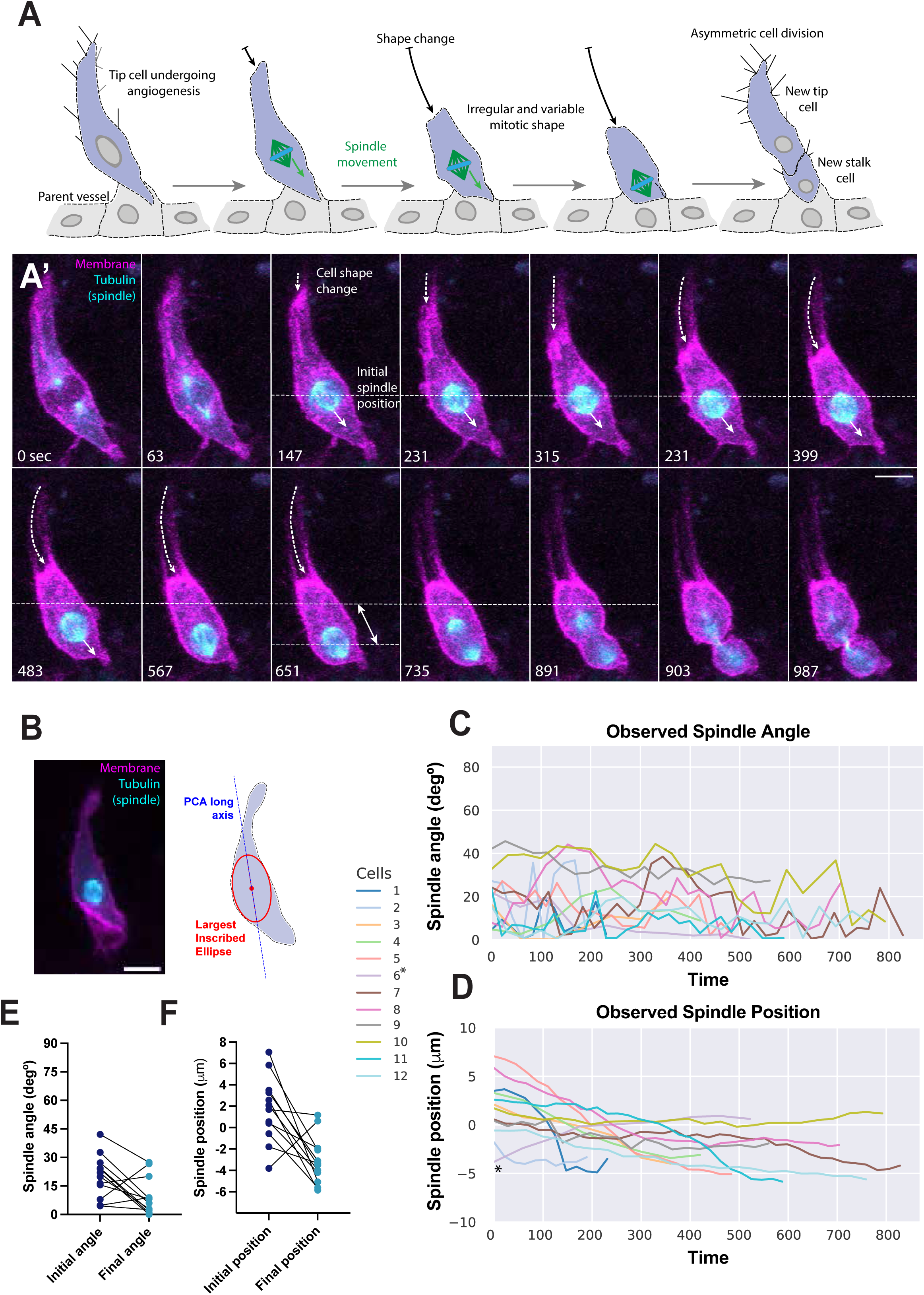
Mitotic spindle dynamics during zebrafish angiogenesis. **(A)** Schematic of a sprouting endothelial cell undergoing mitosis in a zebrafish ISV. Mitotic tip cells undergo limited mitotic rounding, maintaining irregular elongated cell shapes throughout division. Meanwhile, the spindle shifts toward the lower half of the cell, resulting in an off-center division that produces a larger daughter tip cell and a smaller stalk cell. **(A’)** Corresponding microscopy images of the dividing cell shown schematically in **(A)** (Movie S8). Lyn-mCherry (magenta) expressing zebrafish tip endothelial cell with mosaic Tubulin-GFP (cyan) expression undergoing an asymmetric cell division. Arrows indicate change of shape (dashed), movement of mitotic spindle (solid) and total distance spindle has moved (double headed), scale bar 10um. **(B)** Diagram of endothelial cell shape characterization. The cell long axis, determined by principal component analysis, is shown as a blue line. The largest inscribed ellipse is outlined in red, and its center was used as the reference point for position measurements. **(C, D)** Time series of observed spindle angle and position (n = 12, one trace per cell). Each cell is represented by a uniquely colored line. Cell #6, marked with an asterisk, moves upward unlike the other cells (Movie S10). **(E, F)** Initial and final spindle angles and positions for observed (non-simulated) cells. Most of the cells exhibit a downward shift in their spindle position.

Live imaging of mitotic spindle dynamics at high spatial and temporal resolution was performed on twelve cells during ISV growth in zebrafish larvae (as described in (Lovegrove and Herbert 2025)); Cell #11 is shown in Figure 5A (Movie S8). To account for differences in cell size and shape, we determined the area of the largest inscribed ellipse (LIE), a geometry-aware measure of the cell’s usable interior (the “workspace” available for the spindle), capturing both scale and anisotropy (Figure 5B). We used the long axis of this ellipse as a proxy for the long axis of the cell, providing both a basis for spindle angle measurements and a normalization factor to compare the extent of spindle offset between different cells (Figure 5B).

In contrast to the *Drosophila* cells that we have examined, spindle orientation in this system is not very dynamic; the spindle is typically formed at an angle of <45° and remains there through prometaphase, reaching the metaphase-anaphase transition at a final angle of <30° (Figure 5A,C,E). However, the spindle, as previously described does tend to change its position, moving in the proximal direction until it reaches ∼3.5μm below the calculated cell center (Costa et al. 2016) (Figure 5A,D,F).

### Spindle position in simulated endothelial cells is facilitated by asymmetric pulling forces

ISVs develop deep within the zebrafish embryo and are closely associated with many other tissues (muscle, spinal cord, notochord). For this reason, and a lack of fish-specific imaging reagents, the presence/localization of cortical force generators has not been determined in this system. Our computational modeling therefore started with a possibility we considered to be the simplest. Force generators were uniformly distributed around the cortex and astral MTs provided strong pulling and weak pushing force, as used in other cell types above. As before (Figure 4), we also tested pushing only (i.e. the absence of force generators) and pulling-only conditions (Figure 6A). Each cell was modelled individually, including initial spindle angle, position and cell shape, the simulation was then allowed to run altering cell shape at each time step (Figure 5). Forty simulations were undertaken for each cell (Supplemental Figure 5).

**Figure 6:** Computational model predicts that spindle positioning in zebrafish blood vessel cells can be driven by force generators (FGs) localized at cell–cell junctions. **(A)** Schematics of initial three testing scenarios: uniform FGs pulling and pushing, uniform FGs pulling only, and Uniform pushing only. FG localizations are marked in red on the cortex. **(B, C)** Comparison of observed and predicted final spindle angle and position under different simulated conditions. For spindle angle **(B)**, the plotted values are the absolute difference between observed and model predictions. For spindle position **(C)**, the plotted values are the signed difference between observed and model predictions. Boxplots show the 25th, 50th, and 75th percentiles with whiskers spanning the full data range (n = 12 observed cells). Each dot represents the average of 40 simulation runs for a single cell. Conditions are color-coded as follows: blue, combined pulling and pushing with uniformly distributed FGs; red, pulling only with uniformly distributed FGs; green, pushing only without FGs; purple, combined pulling and pushing with FGs localized to cell–cell junctions; pink, pulling only with FGs localized to cell–cell junctions. Statistical significance was assessed using the paired Wilcoxon signed-rank test; *p < 0.05, **p < 0.01, ***p < 0.001, ****p < 0.0001. **(D)** Left: cell–cell junctions at the base of a branching endothelial cell, which may serve as polarity cues biasing the spindle downward. Lyn-mCherry (magenta) expressing zebrafish tip endothelial cell with mosaic Tubulin-GFP (cyan) undergoing an asymmetric cell division. Cell–cell junctions are marked by ZO-1 (white). **(E)** Schematic of a cell with FGs restricted to cell–cell junctions, giving rise to two tested conditions—junction pulling and pushing, and junction pulling only. **(F,G)** Paired plots comparing observed values (cyan) of final spindle angle **(F)** and position **(G)** with predictions (purple) from the combined pulling and pushing condition with junctional FG localization. The line marked with an asterisk indicates an outlier cell, in which the spindle exhibited an atypical upward shift (Movie S10).

Spindle angles were close to observation (an average difference of ∼10°) in both of the conditions that incorporated pushing, whereas spindle orientation was unpredictable in the uniform pulling-only condition (Figure 6B). A possibility suggested by earlier work is that pushing MTs provide a readout that translates cell shape into spindle orientation, and our modeling results agree with that possibility. However, the spindle failed to reach its observed final position – proximal to the rear end of the cell – in all cases (Figure 6C). This result suggested an asymmetry in pulling, as observed in the *C. elegans* embryo (modeled in Figure 3).

In mammalian epithelial cells, the spindle orienting factor LGN (homologous to *Drosophila* Pins and *C. elegans* GPR1/2) localizes to cell-cell junctions through direct interaction with E-Cadherin (Gloerich et al. 2017). This interaction can also occur between LGN and vascular specific VE-Cadherin (Monster et al. 2024). We therefore considered the possibility that force generators in the tip cell are located at cell-cell junctions. Tip cells share a junction only with a single adjacent cell (belonging to either the parent blood vessel or trailing stalk cell), as visualized by staining for the tight junction component ZO-1 (Figure 6D). Junctional proteins are therefore limited to the proximal region of tip cell. This presents two more conditions for our model: junctional pulling with pushing and junctional pulling alone (Figure 6E). Spindle orientation was not reliable in the latter condition (Figure 6B). Moreover, the final spindle position was even further towards the proximal end of the cell than the observed positions by an average of ∼2μm (Figure 6C). In agreement with results presented in Figure 4, this indicates that MT pushing forces provide a buffer that prevents spindle poles from getting too close to the cortex. The junctional pulling and pushing condition provided the most accurate recapitulation of both spindle angle and position (Figure 6 B,D,F,G). This was true across a range of astral MT and force generator numbers (Supplemental Figure 6A,B).

This work demonstrates that pushing forces generated by astral MTs alone, coupled with the elongated cell shape maintained by these cells during mitosis, are sufficient to correctly orient mitotic spindles in this system. However, an asymmetric pulling force (potentially localised via cell-cell junctions) is additionally required to correctly position the spindle. Therefore endothelial tip cells appear to rely on a specific combination of astral MT based pushing and pulling forces, along with elongated cell shapes, to generate their specific spindle dynamics.

### Limitations of this study

While our model successfully captures essential features of spindle dynamics across multiple systems, there are simplifications that should be considered. Our model is restricted to a 2D midplane, which neglects interactions between out-of-plane MTs and FGs, that can lead to off-axis forces and torques. Second, we assumed ideal Euler beam behavior for MT buckling and modeled cortical force generators as static entities with fixed pulling strengths, whereas real MTs may deviate from this behavior due to bundling effects, structural imperfections and regulatory proteins, potentially altering buckling behavior and force predictions. Moreover, it is possible that astral MTs pushing against the cortex can slip along the surface, depending on the angle of incidence. Such behavior would alter the spatial organization of MTs and the distribution of forces. Additionally, the assumption of a rigid, non-deformable cortex may be oversimplified for cells undergoing significant shape changes during division.

## DISCUSSION

Our computational modeling reveals that MT pushing forces play critical but context-dependent roles in mitotic spindle dynamics, working synergistically with cortical pulling mechanisms to ensure robust orientation and positioning across diverse cell types and geometries. While previous work has established cortical pulling as the primary mechanism for spindle orientation, our results reveal three distinct ways that pushing forces contribute to spindle dynamics. Firstly, pushing MTs act as geometric sensors that translate cell shape into spindle behavior. Because pushing force is inversely proportional to MT length, the spindle receives continuous feedback about its position relative to the cell cortex. This mechanism is particularly evident in elongated endothelial cells, where shape-dependent pushing forces alone are sufficient to achieve appropriate spindle orientation. Secondly, pushing forces provide a “buffering” function that prevents spindle poles from approaching the cortex too closely. When spindles rely solely on cortical pulling, they can become stalled in suboptimal orientations, with one pole trapped near cortical force generators while the other is pulled away from the opposing side. Pushing MTs counteract this by generating repulsive forces that scale with proximity to the cortex, effectively creating a dynamic equilibrium that maintains spindle mobility. This buffering effect is most critical at high initial spindle angles, where the risk of stalling is greatest. Thirdly, pushing forces promote overall spindle dynamicity by driving rotational movements that increase the likelihood of productive encounters between spindle poles and cortical force generators. While pushing forces themselves are non-directional, they accelerate the search process by which pulling mechanisms find their targets, thereby reducing the time required for proper alignment.

The relative importance of pushing and pulling mechanisms varies systematically across cell types in ways that reflect their geometric and functional requirements. In symmetrically dividing cells with regular geometries, such as *Drosophila* follicle cells, cortical pulling provides the primary directional cue while pushing forces enhance efficiency and prevent stalling. The uniform distribution of astral MTs in these cells means that pushing forces are approximately isotropic, making them unsuitable for establishing polarity but ideal for maintaining spindle mobility.

In contrast, asymmetrically dividing cells like *C. elegans* embryos and zebrafish endothelial tip cells require precise control over spindle position to generate daughters of appropriate size and fate. Here, asymmetric distributions of cortical force generators create the positional bias necessary for off-center spindle placement, while pushing forces ensure that spindles can still move dynamically despite being displaced from the geometric center. The elongated morphology of dividing endothelial cells represents an extreme case where cell shape itself, combined with isotropic pushing, is sufficient to orient the spindle correctly.

Our modeling results suggest a role for cortical force generators in zebrafish tip cell division. While these have not been explored directly in zebrafish, LGN is expressed in Human Umbilical Vein Endothelial Cells (Wright et al. 2015), which undergo angiogenic sprouting behavior in culture. Notably, LGN is not required for spindle orientation in these cells, but it is required for proper endothelial sprouting and branching (Wright et al. 2015). These results are consistent with our model predictions that endothelial cells can achieve spindle orientation through shape-dependent pushing forces, while still requiring cortical machinery for proper positioning and function.

This work establishes a new framework for understanding mitotic spindle control that emphasizes the cooperative nature of pushing and pulling mechanisms. Rather than competing alternatives, these forces function as complementary components, providing both robustness and precision. This dual-mechanism framework resolves experimental contradictions and suggests that therapeutic interventions should consider the full spectrum of underlying mechanisms. As our understanding continues to evolve, similar computational approaches will prove valuable for dissecting the mechanical basis of cellular organization across diverse biological systems.

## Materials and Methods

### Computational Model

We defined four key components: the mitotic spindle, astral MTs, the cell cortex, and cortical force generators (FGs). The mitotic spindle is modeled as a rigid rhombus with a long diagonal (2r) and a short diagonal (2w). The long diagonal represents the spindle length, while the short diagonal represents spindle width, and both are kept constant throughout the simulation, reflecting the observed stability of spindle dimensions during metaphase in most cell types (Dumont and Mitchison 2009). The spindle has an initial angle θ, with centrosomes positioned at the ends of the long diagonal. Spindle position and angle are updated iteratively at each time step based on the net force and torque arising from all astral MTs.

Astral MTs nucleate from the centrosomes within a restricted anterior β angle (Supplemental Table 1), under the assumption that MTs directed toward the metaphase plate are obstructed and cannot reach the cortex. MTs grow with velocity v_g_ and shrink with velocity v_s_. Their initial phase—growing or shrinking—is set randomly, and their initial lengths are also randomized to reflect prior nucleation from duplicated centrosomes. Once MTs grow long enough to reach the cortex, they can either (1) push against it, generating a pushing force, or (2) bind to an available FG within range and pull. These events are modeled stochastically. Dynamic instability (rescue and catastrophe) as well as FG binding and unbinding are implemented using a Poisson process: at each time step, a pseudorandom number (random(0,1)) is sampled and compared to the event probability 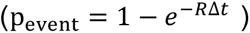 derived from the corresponding rate constant (*R*). The random event is going to occur if p_event_ < random(0,1). This stochastic implementation was adapted from previous spindle modeling frameworks (Civelekoglu-Scholey et al. 2006; Zhu et al. 2010).

Cortical FGs are distributed according to cell-type–specific patterns or experimental conditions. Each FG can bind at most one astral MT. If an MT contacts the cortex within distance *d*_*bind*_ of a free FG, binding occurs with probability 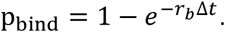 Once bound, the FG exerts a constant pulling force (approximated by the dynein stall force) directed along the line connecting the MT and FG. Bound MTs can detach with probability 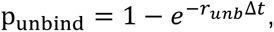 in which case pulling will stop.

For follicle cells, neuroblasts, and C. elegans embryos, we distributed a fixed number of force generators (FGs). In contrast, for endothelial cells, we used an FG density–based approach, so that the total number scaled with cell size. This ensured comparable conditions across cells of varying dimensions, while still allowing us to localize FGs to specific cortical regions depending on the condition being tested.

MTs that reach the cortex while in the growing state and without binding to an FG exert pushing forces via polymerization. These are modeled using Euler buckling mechanics, 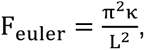 where κ is a MT bending rigidity and L is its length. Pushing force per MT is capped at a maximum value of 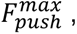 which we refer to in this work as the “pushing strength”. This value is consistent with experimental estimates (Dogterom and Yurke 1997), so pushing force becomes 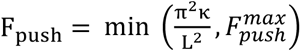 as in (Li and Jiang 2017). To investigate its impact on spindle dynamics, we varied the pushing strength across a physiologically reasonable range of 0.5–10 pN. Unless otherwise specified, both pulling and pushing strengths were kept at 5 pN.

The cytoplasm is modeled as a viscous medium (viscosity η) in the low Reynolds number regime, where spindle motion follows a Langevin-type description. The effective drag on the spindle is approximated as that of a sphere, 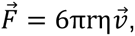 and we neglect explicit drag on individual astral MTs (Kimura and Onami 2005). While this approximation may underestimate the absolute drag, it primarily acts to rescale dynamical time scales compared to full viscous drag description.

To find the net force acting on the spindle, we sum the forces 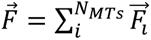 over all individual astral MTs. We assume that the spindle only rotates around its center of mass. To find net torque acting on the spindle, we calculate total force acting on each spindle pole 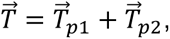 where 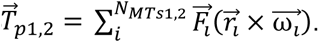 From total torque 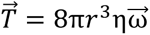 we can calculate angular velocity of the spindle.

The total force gives the translational position change.

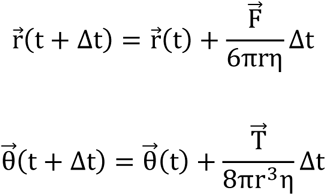

Because of stochastic variability, individual runs yield diverse outcomes. To obtain robust results, we performed multiple independent simulations per condition. The number of runs was increased until averages stabilized (Lee et al. 2015).

We used a 0.05 s simulation time step because it provides a good balance between computational efficiency and temporal resolution. Larger time steps can result in the spindle moving unrealistically far in a single update, leading to inaccurate representations of force balance and spatial regulation. All parameter values are listed in Supplemental Table 1.

In our model, the spindle can stall when any attempted movement—either translational or rotational—would result in a violation of the cell’s geometric boundary. Specifically, this occurs when the predicted displacement places spindle outside the permissible actin cortex, which is modeled as a rigid, non-deformable wall. Such situations typically arise when the spindle is near the cell periphery and the existing pushing or pulling forces are insufficient to reposition it without crossing the boundary. However, due to the stochastic nature of astral MT dynamics, changes in the force profile may occasionally rescue these stuck spindles by enabling a permissible displacement.

### Live imaging of mitotic spindles in zebrafish ISVs

Embryos and adults were maintained under standard laboratory conditions as described previously (1) and experiments were approved by the University of Manchester Ethical Review Board and performed according to UK Home Office regulations. For experiments, embryos were not selected for gender and no statistical method was used to predetermine sample size for experimental groups.

To generate mosaic expression of tubulin-GFP 32pg of pTol2 Fli:GFP-alpha-tubulin (2) was co-injected with 32pg Tol2 mRNA into one-cell stage Tg(kdrl:ras-mCherry)s896 (3) zebrafish embryos. Embryos were then grown to approximately 24hpf at 28°C, dechorionated and mounted in 1% low-melting agarose (containing 0.1% tricane) in glass-bottom dishes. They were continually perfused with embryo water supplemented with 0.0045% 1-phenyl-2-thiourea and 0.1% tricane using an in-line solution heater and heated stage. For further details see Lovegrove and Herbert, 2024 (4). Embryos were imaged using a 20x or 40x dipping objective on either a Zeiss 700 or 980 Airyscan confocal microscope. Z-stacks were acquired every 20sec-1min, with slices set 2um apart.

Staining of embryos was carried out by fixing ∼24hpf embryos in 4% PFA overnight at 4°C. Then washed 3 times in 0.1% PBST for 5mins and permeablised in PBS + 2% Triton-100 for 1.5hrs at room temperature (RT). Blocked in PBS with 1%BSA, 0.1% Triton-X100 and 2% heat treated lamb serum for a least an hour at RT. ZO-1 primary antibody (Invitrogen 33-9100) was diluted at 1:250 in block and incubated overnight at 4°C. Finally, they were washed in block at RT for at least 8hrs before incubating them with secondary antibody (Invitrogen A11004) at 1:200 in block overnight at 4°C. They were then washed, mounted and imaged as above.

### Implementation of cell shape change over time

The images of individual cells were obtained from live-cell imaging experiments and stored as binary mask images for both the cell boundary and the mitotic spindle. Image processing and geometric quantification were performed in Python using OpenCV (cv2) together with numerical and geometric analysis libraries (numpy, shapely, scipy). For each cell, the boundary contour was extracted from the cell mask at all imaged time points and stored, providing a discrete series of cell shapes. Because the shape updates are only available at these discrete time points, changes in cell geometry during simulations would also be discrete. To approximate a continuous evolution of cell shape, we applied linear interpolation between consecutive time points, generating an interpolated cell contour every 1 second. While the true intermediate shapes are unknown, this approach provides a smooth approximation for simulations, allowing the cell boundary to evolve continuously over time.

### Quantification of cell geometry and long-axis determination in Zebrafish

To ensure a consistent reference frame, all cell contours were translated such that the origin of the coordinate system corresponded to the experimentally measured initial spindle center position. This translation was achieved by detecting the spindle contour from the corresponding spindle mask image and subtracting the spindle centroid coordinates from all cell boundary coordinates.

The cell’s long axis was defined as the major axis of the *largest inscribed ellipse* that could be fully contained within the observed cell contour. To identify this ellipse, we employed an iterative fitting algorithm that systematically varied the ellipse center position, semi-major and semi-minor axes lengths, and orientation, while ensuring that the generated ellipse was entirely contained within the cell polygon. The ellipse with the maximal area was selected, and its orientation (in the range 0–180°) was taken as the long axis angle. The center of this fitted ellipse served as a geometric reference point for subsequent measurements.

Distances between experimental spindle positions and the fitted ellipse center were computed in two forms: 1) Euclidean displacement — the straight-line distance between the spindle position and ellipse center; 2) Projected displacement along the long axis — the component of the displacement vector aligned with the ellipse major axis.

These metrics provided a standardized way to quantify spindle positioning relative to intrinsic cell geometry. All coordinates and distances were reported in micrometers.

## Acknowledgments

This work was supported by NSF CAREER award 2042280 (PI: Bergstralh), British Heart Foundation Accelerator Award AA/18/4/34221 (PI: Lovegrove), EPSRC award EP/W024144/1 (PI: Fletcher), and BBSRC award BB/Y514020/1 (PI: Fletcher). Data were partially acquired at the University of Missouri Advanced Light Microscopy Core facility.

## Data availability

The authors affirm that all data necessary for verifying the conclusions are included in the text, figures, and tables.

---

**Supplemental Figure 1:**
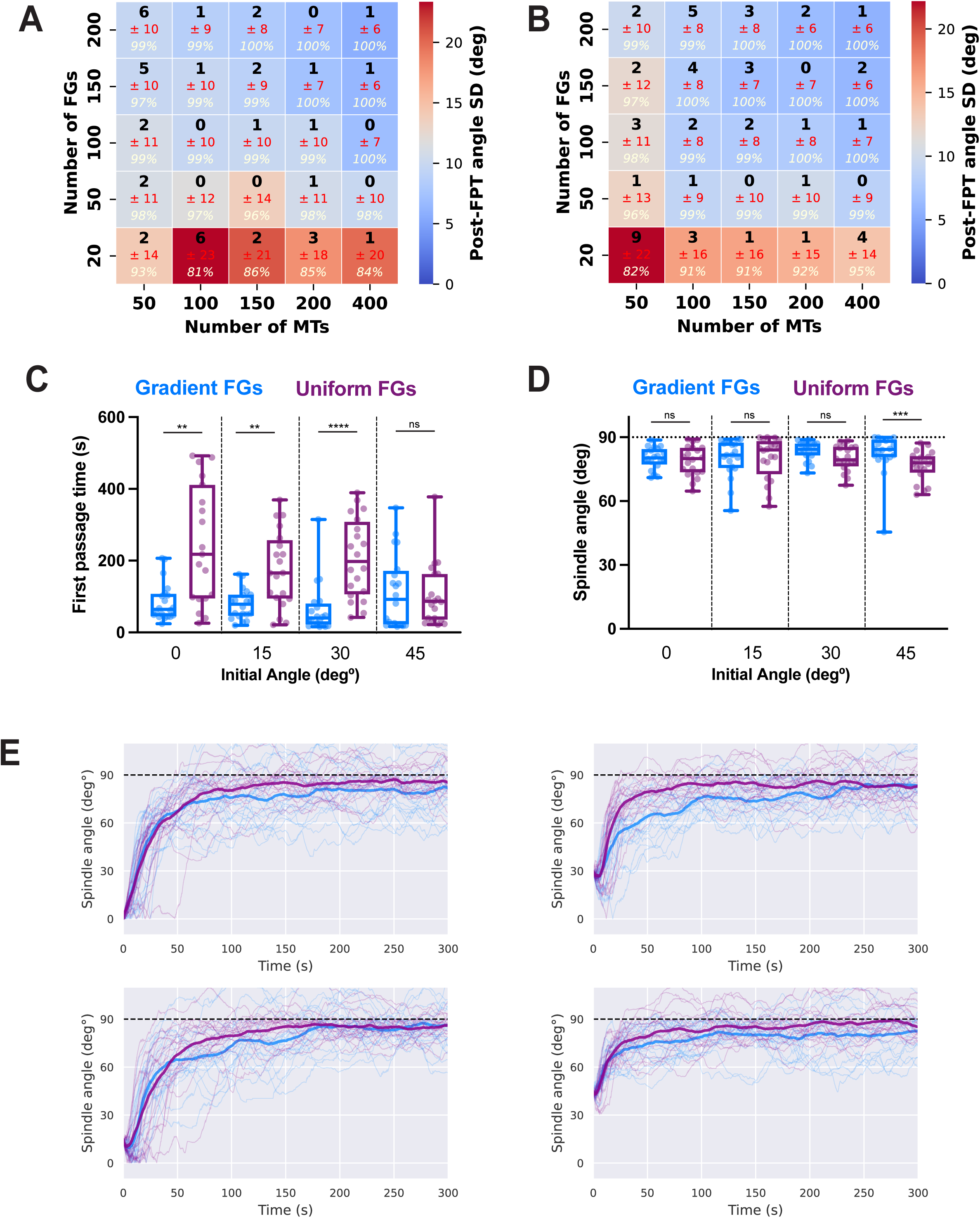
Effects of FG localization patterns and force balance on spindle orientation. Heatmaps of post-FPT spindle angle stability as a function of number of MTs and number of FGs for follicle cells **(A)** and neuroblasts **(B)** in the combined pulling and pushing condition. For each parameter pair, mean post-FPT angle is shown in bold, the standard deviation in red, and the percentage of time the spindle remained within the target alignment window in italics. Heatmap color encodes spindle angle standard deviation, with red indicating the largest variation and blue the smallest. Both pulling and pushing strengths were fixed at 5 pN. **(C, D)** First passage time and final spindle angle under different FG localization patterns and initial spindle angles. Spindles with non-uniformly localized FGs orient more quickly **(C)**, but localization patterns have little impact on final spindle angle when comparing uniform-like versus gradient-like distributions **(D)**. Boxplots show the 25th, 50th, and 75th percentiles with whiskers spanning the full range (n = 20 runs). Conditions are color-coded as follows: blue, combined pulling and pushing with gradient-distributed FGs; dark purple, combined pulling and pushing with uniformly distributed FGs. Statistical significance was assessed using the Mann-Whitney test; *p < 0.05, **p < 0.01, ***p < 0.001, ****p < 0.0001. **(E)** Spindle angle over time in simulated neuroblast cells. Initial spindle angles were 0°, 15°, 30°, and 45° (top to bottom, left to right). Lines are color-coded as in **(C)**; solid lines show the mean, and thin lines represent individual runs.

**Supplemental Figure 2:**
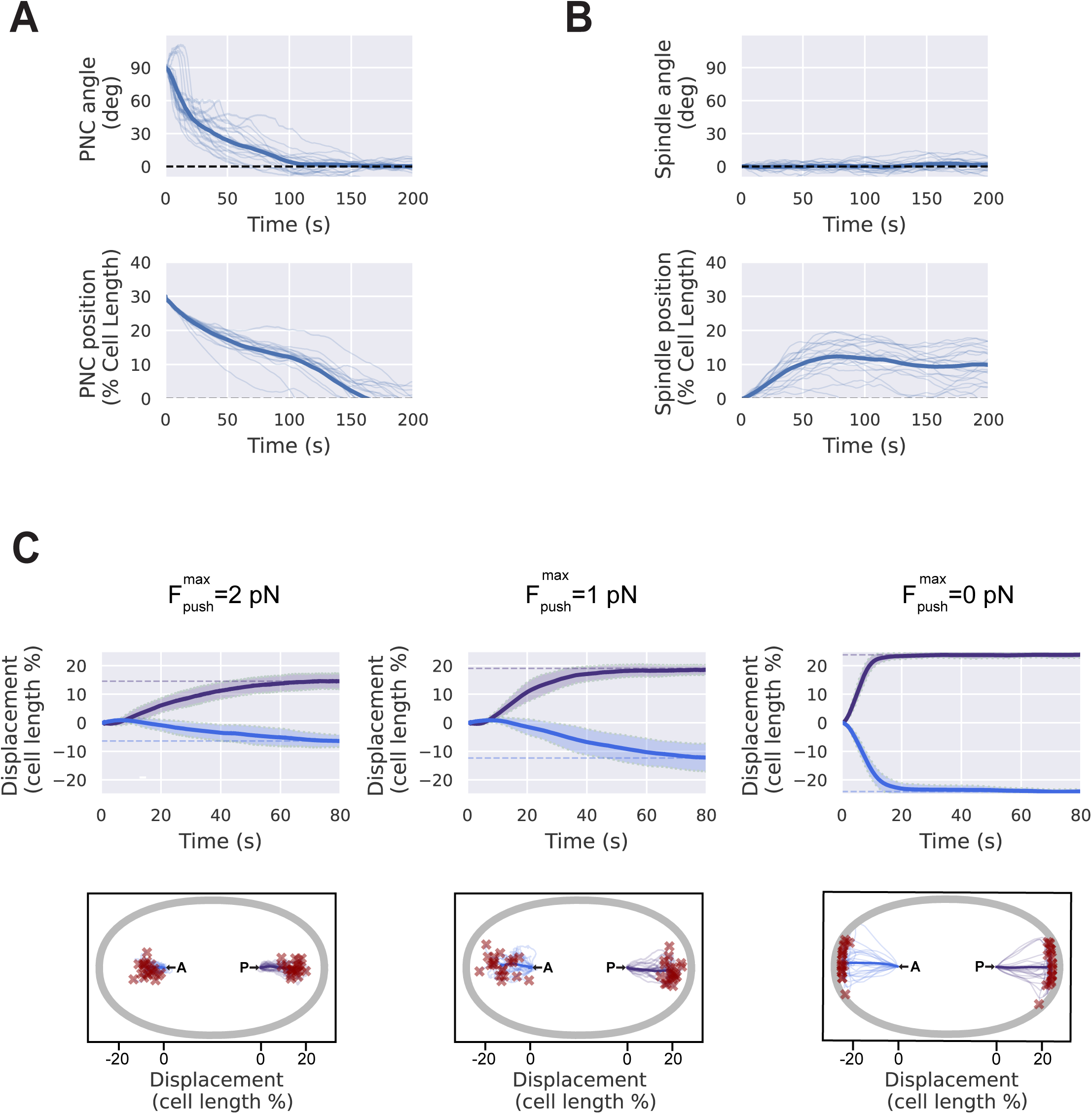
Individual simulation trajectories for C. elegans model validation. **(A)** Pronuclear complex (PNC) migration. Time series of PNC angle (top) and position (bottom). All 20 individual simulation runs are shown as separate traces. **(B)** Posterior spindle displacement. Time series of spindle angle (top) and position (bottom). All 20 individual runs are shown as separate traces. **(C)** Simulated spindle severing, shown left to right, under cortical pulling with decreasing pushing strengths (2 pN, 1 pN, and 0 pN). Top: Position versus time plots for posterior (purple) and anterior (blue) poles following severing of a centered spindle (L = 22 μm) at t = 0 s. Solid lines represent the mean of n = 20 runs; dashed horizontal lines are initial positions for both poles. Bottom: Scatterplot of spindle pole trajectories (color coded as before) and final positions (red X marks). All 20 individual runs are shown as separate traces. *N*_*MT*_ =1000 and *N*_*FG*_ = 200.

**Supplemental Figure 3:**
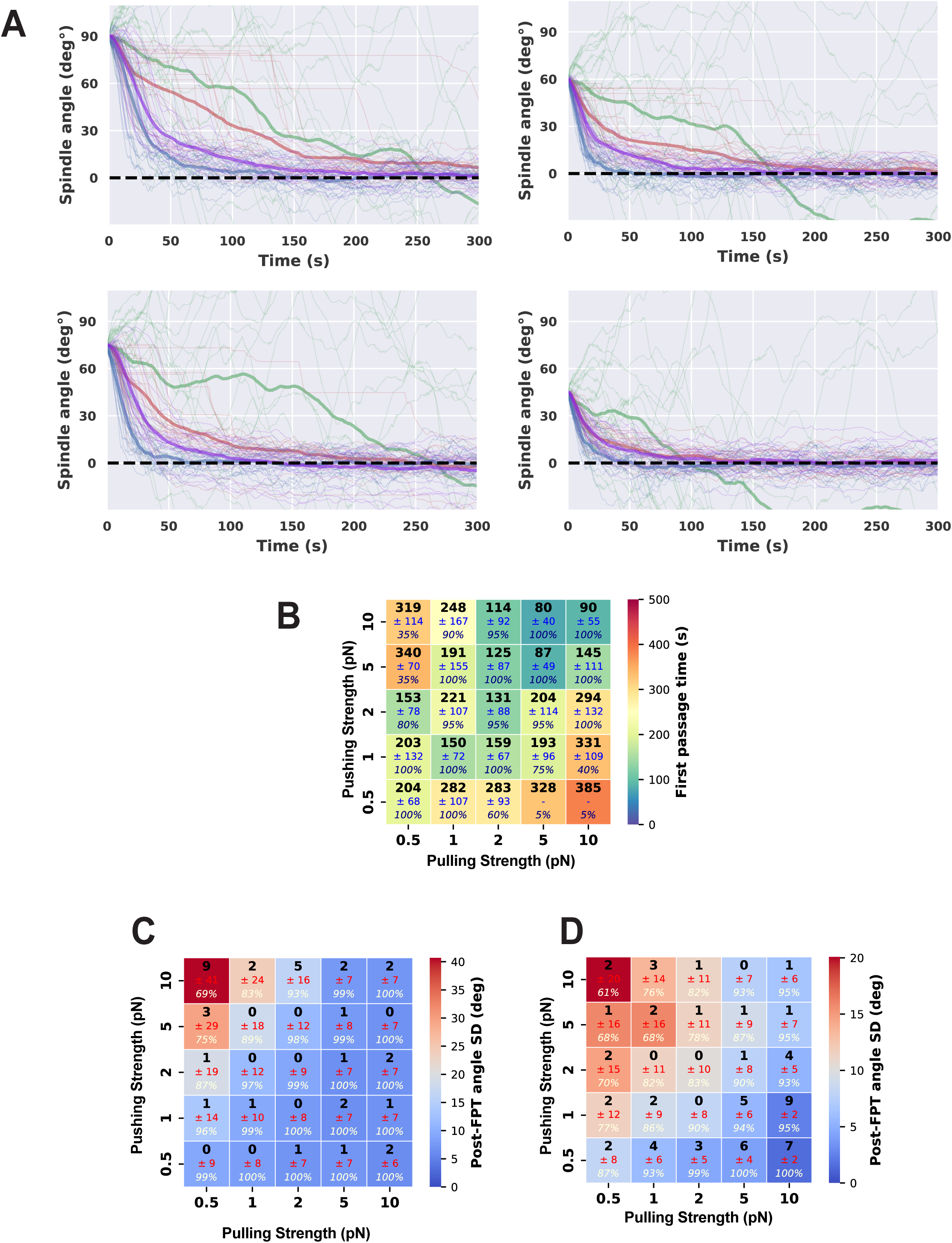
Additional analysis of pushing forces in follicle cell simulations. **(A)** Spindle angle over time for simulated follicle cells with initial angles of 90°, 75°, 60°, and 45° (top to bottom). Four conditions are shown: combined pulling and pushing (blue), pulling only (red), pushing only (green), and pinned pulling only (purple). Solid lines represent the mean of n = 20 simulations, while thin lines show the individual runs. **(B)** Dependence of spindle orientation speed on MT pulling and pushing strength in the combined condition (*N*_*MT*_ =200 and *N*_*FG*_ = 100) in neuroblasts. For each parameter combination, the average first passage time (FPT, n = 20 runs) is plotted in bold black, with standard deviation in navy and percent of successful orientations in italics. **(C, D)** Heatmaps showing how spindle stability depends on variations in pulling and pushing strength under the combined pulling and pushing condition in follicle cells **(C)** and neuroblasts **(D)**. Stability is quantified by post-FPT spindle angle standard deviation (bold black), with standard deviation in navy and the percentage of time the spindle remained within the target alignment window in italics. Heatmap color encodes spindle angle standard deviation, with red indicating the largest variation and blue the smallest. *N*_*MT*_ =100 and *N*_*FG*_ = 200.

**Supplemental Figure 4:**
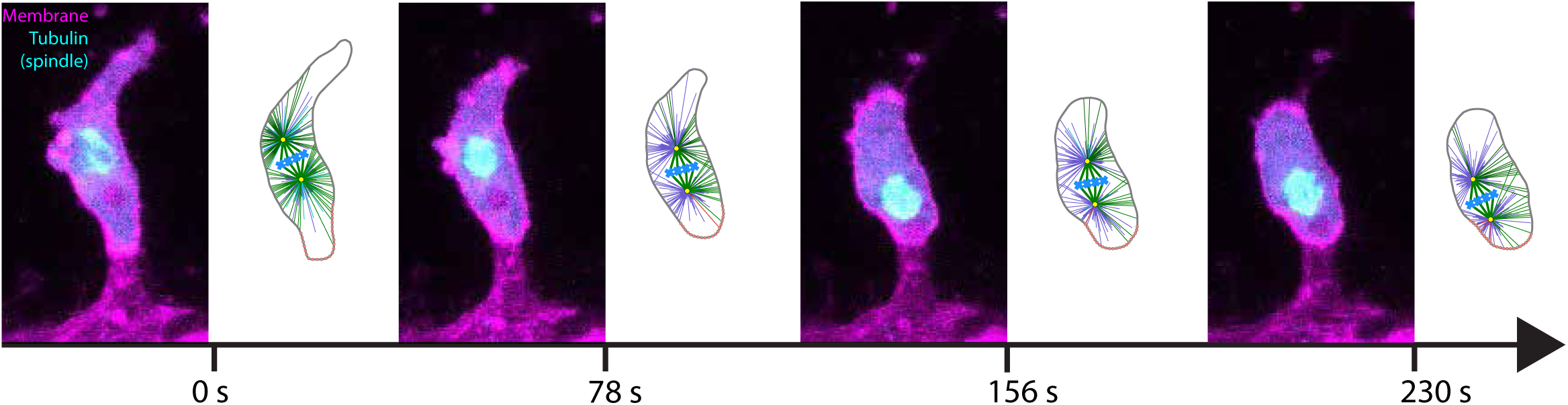
Microscopy images alongside representative simulation snapshots at corresponding time points, illustrating how the model reproduces observed cell shape changes during division.

**Supplemental Figure 5:**
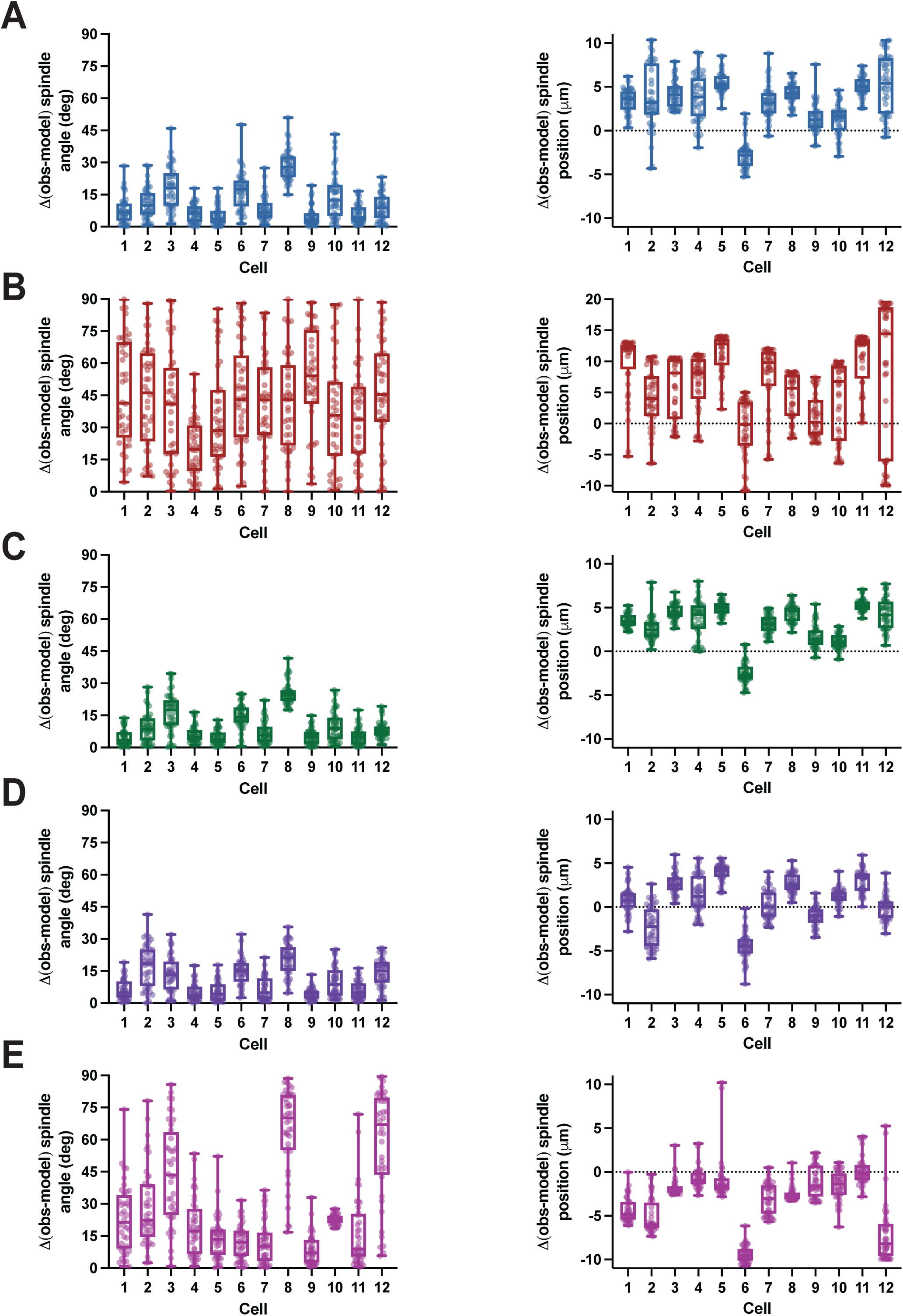
Detailed comparison of observed and predicted final spindle angle and position under different simulated conditions. This figure expands on the summary data shown in Figure 6B–C by presenting results for each cell and each simulated condition separately. For spindle angle (left column), values represent the absolute difference between observed and predicted outcomes. For spindle position (right column), values represent the signed difference between observed and predicted outcomes. Boxplots display the 25th, 50th, and 75th percentiles, with whiskers spanning the full data range (n = 40 simulation runs per cell). Simulation conditions are color-coded as follows: blue (A), combined pulling and pushing with uniformly distributed FGs; red (B), pulling only with uniformly distributed FGs; green (C), pushing only without FGs; purple (D), combined pulling and pushing with FGs localized to cell–cell junctions; pink (E), pulling only with FGs localized to cell–cell junctions.

**Supplemental Figure 6.** Junctional pulling and pushing robustly predict spindle angle and position across parameter settings. Comparison of observed and predicted final spindle angle (**A**) and position (**B**) under different simulated conditions across multiple parameter pairs (*N*_*MT*_ - *N*_*FG*_). For spindle angle (left), values represent the absolute difference between observed and model predictions. For spindle position (right), values represent the signed difference between observed and model predictions. Boxplots show the 25th, 50th, and 75th percentiles with whiskers spanning the full range (n = 12 observed cells). Each dot corresponds to the average of 40 simulation runs for a single cell. Conditions are color-coded as follows: blue, combined pulling and pushing with uniformly distributed FGs; red, pulling only with uniformly distributed FGs; green, pushing only without FGs; purple, combined pulling and pushing with FGs localized to cell–cell junctions; pink, pulling only with FGs localized to cell–cell junctions.

**Movie S1:** Spindle orientation in the simulated follicular epithelium cell from Figure 2A.

**Movie S2:** Spindle orientation in the simulated neuroblast from Figure 2A’.

**Movie S3:** Simulated pronuclear complex (PNC) migration in the *C. elegans* embryo from Figure 3A’’.

**Movie S4:** Spindle positioning in the *C. elegans* embryo from Figure 3B’’.

**Movie S5:** Simulated spindle severing in the *C. elegans* embryo with cortical pulling and pushing (related to Figure 3C”).

**Movie S6:** Simulated spindle severing in the *C. elegans* embryo with cortical pulling only (related to Supplemental Figure 2C).

**Movie S7:** Representative simulation of a “stalled” spindle.

**Movie S8:** Dividing zebrafish endothelial cell corresponding to Figure 5A’.

**Movie S9:** Dividing zebrafish endothelial cell corresponding to Supplemental Figure 4.

**Movie S10:** The outlier cell, in which the spindle exhibited an atypical upward shift.

**Movie S11:** Simulation of uniform pulling and pushing (UF pulling + pushing).

**Movie S12:** Simulation of uniform pulling only (UF pulling). **Movie S13:** Simulation of uniform pushing only (UF pushing). **Movie S14:** Simulation of junction-localized pulling and pushing.

**Movie S15:** Simulation of junction-localized pulling only.

**Supplemental Table 1.**

|  | Parameter | Simulation Value | Description/ Reference |
| --- | --- | --- | --- |
| **Cell** |  |  |  |
| | Cytosol viscosity ($\gamma$) | 1 Pa·s | (Kimura and Onami 2005) |
| | Cell length ($L_{cell}$) and width ($W_{cell}$): | | |
| | *C. elegans* | 50 μm, 30 μm | (Kimura and Onami 2005) |
| | Neuroblast | 10 μm, 10 μm | Based on observations from (Cabernard and Doe 2009) |
| | Follicle cell | 10 μm, 10 μm | Based on observations from Neville et al. (2023) |
|  | Zebrafish | - | Cell contours were taken from microscopy images. |
| **Spindle** |  |  |  |
| | Spindle length ($L_{cell}$) and width ($W_{cell}$): | | |
| | *C. elegans* (PNC) | 10 μm, 10 μm | (Kimura and Onami 2005) |
| | *C. elegans* (positioning) | 12 μm, 10 μm | Adapted from Figure 1 (Pécéréaux et al. 2016) |
| | *C. elegans* (laser ablation) | 22 μm, 10 μm | Estimate of final spindle length |
| | Neuroblast | 8 μm, 5 μm | Based on observations from (Cabernard and Doe 2009) |
| | Follicle cell | 8 μm, 5 μm | Based on observations from Neville et al. (2023) |
|  | Zebrafish | - | Spindle contours were taken from microscopy images. |
| **Microtubules** |  |  |  |
|  | Catastrophe rate | 0.021 s<sup>-1</sup><br>0.014 s<sup>-1</sup> (*C. elegans*) | (Rogers et al. 2002)<br>(Kimura and Onami 2005) |
|  | Rescue rate | 0.029 s<sup>-1</sup><br>0.044 s<sup>-1</sup> (*C. Elegans*) | (Rogers et al. 2002)<br>9/23/2025 12:18:00 AM |
| | Growth rate ($v_g$) | 0.13 μm/s | (Burakov et al. 2003) |
| | Shrink rate ($v_s$) | 0.27 μm/s | (Burakov et al. 2003) |
| | Flexural rigidity ($\kappa$) | 20 pN·μm<sup>2</sup> | (Nédélec 2002) |
| | Number of MTs ($N_{MT}$) | 50-1000 | We varied this number throughout this work. |
| **Forces** |  |  |  |
| | Dynein stall force ($F_{pull}$) | 0.5-10 pN | (Toba et al. 2006) |
| | Dynein binding rate ($r_b$) | 0.03 s<sup>-1</sup> | (Li and Jiang 2017) |
| | Dynein unbinding rate ($r_{unb}$) | 0.02 s<sup>-1</sup> | (Li and Jiang 2017) |
| | Polymerization stall force ($F_{push}^{max}$) | 0.5-10 pN | (Dogterom and Yurke 1997) |
| **Model parameters** |  |  |  |
| | Time step ($\Delta t$) | 0.05 s | This interval is short enough to capture the fast dynamics of microtubule interactions and spindle movements without introducing numerical instability or skipping over critical transitions. |
| | MTs angular spread ($\beta$) | 270°<br>180° (*C. elegans* PNC) | MT angular spread excludes directions toward metaphase plate due to obstruction by chromosomes and pro nuclei. |

